# Contrasting mutation patterns in haploid and diploid cells from two yeast species

**DOI:** 10.1101/2025.09.30.679554

**Authors:** Kevin Bao, Rutuja Gupte, Neil Braker, Nathaniel P. Sharp

## Abstract

There is significant variation in the rate and spectrum of spontaneous mutations among taxa. How this variation is shaped by natural selection remains a subject of debate. The drift barrier hypothesis proposes that selection generally favors lower mutation rates due to the risk of deleterious mutations but acts less effectively against weak mutator alleles in smaller populations, allowing the mutation rate to increase due to genetic drift. Given this model, we propose that mutation rates may also be elevated in cell types that appear rarely in a population, where DNA replication and repair processes are subject to selection less often. We can begin to test this prediction in yeast species, some of which can be grown in either a haploid or diploid cell state. Existing data on the budding yeast *Saccharomyces cerevisiae* support this prediction, with a higher mutation rate observed in haploids, which is the rare cell type in natural populations. However, this pattern could also appear if haploidy is inherently mutagenic, regardless of the dominant cell type. To test these alternatives, we conducted a mutation accumulation experiment with haploid and diploid cells of the fission yeast *Schizosaccharomyces pombe*, in which diploidy is the rare cell type. In this species, we found a higher mutation rate in diploids, consistent with our prediction. In both species, the spectrum of mutations is also influenced by ploidy state. Our findings suggest that limits to selection on mutation may be evident as variation within species.

**Significance:** Spontaneous mutation rates vary among organisms. Natural selection may act to reduce mutation rates; if so, we would expect mutation rates to be elevated in cell types where natural selection has historically had less opportunity to act. We studied mutation patterns in two yeast species when grown in the haploid or diploid form. For the typically diploid species, the mutation rate was higher in the haploid form. For the typically haploid species, the mutation rate was higher in the diploid form. The observation that mutation rates increase when selection is ineffective indicates that selection usually acts to reduce mutation rates.

## Introduction

Spontaneous mutations provide material for adaptation but often have deleterious fitness effects. The relatively high fidelity of genome transmission suggests that natural selection has generally favored lower mutation rates over the course of evolution. However, there is considerable variation in the rate and spectrum of mutations across the tree of life. A possible explanation for this variation is that selection favors different mutation rates in different organisms, with an “optimal” mutation rate determined by the cost of replication fidelity and the expected frequencies of beneficial and deleterious mutations (Sniegowski et al. 2000; Baer et al. 2007). Alternatively, selection could generally favor lower mutation rates in all organisms, with mutation rate diversity reflecting variation among populations in the effectiveness of selection against mutator alleles, determined by their realized fitness effects and the effective population size, *N_e_*. Alleles causing small increases in the mutation rate (weak mutator alleles) will likely cause small reductions in fitness, making them less subject to purifying selection in populations with lower *N_e_*. This is known as the drift barrier hypothesis for mutation rate evolution (Lynch 2008, 2011).

A key source of empirical support for the drift barrier hypothesis is a negative correlation between estimates of mutation rate and *N_e_* across taxa (Lynch 2010; Lynch et al. 2016), but this conclusion is disputed. One issue is that mutation rates are typically needed to estimate *N_e_*, which might contribute to a negative correlation between mutation rate and *N_e_* due to sampling error alone (Sung et al. 2012; Wang et al. 2020). There are additional potential challenges in estimating *N_e_*, including the effects of linked selection (Waples et al. 2016; Kern and Hahn 2018; Johri et al. 2021; Latrille et al. 2021), and selection on synonymous variants (Chamary et al. 2006; Lebeuf-Taylor et al. 2019; Bailey et al. 2021; Shen et al. 2022; Zhang and Qian 2025). Additionally, some experiments suggest that relaxed selection leads to the evolution of increased mutation rates, as expected under the drift-barrier hypothesis (Saxena et al. 2018; Wei et al. 2022), but others do not (Liu and Zhang 2021a). Additional approaches to testing the drift barrier hypothesis would be valuable.

Notably, the drift-barrier hypothesis is also expected to apply to the relative efficacy of alternative DNA repair pathways within organisms. If selection has less opportunity to act on a given DNA repair pathway, because it is employed infrequently, alleles degrading that pathway will be subject to less effective selection and can therefore reach high frequencies in the population. In support of this idea, there is evidence that polymerases used infrequently to replicate small amounts of DNA are particularly error-prone (Lynch 2008). Similarly, in yeast and mammals, error rates are higher for those proofreading and mismatch repair mechanisms that are active “downstream” in the DNA replication process, which impact fewer nucleotides than the initial polymerization machinery (Lynch 2011). Here, we sought to explore whether this logic also applies to alternative cell types––haploids and diploids––in unicellular fungi.

In the budding yeast *S. cerevisiae*, diploid cells largely divide asexually, but meiosis can be induced by a lack of nitrogen; the resulting haploids are of two mating types and will readily mate with a nearby haploid cell of the opposite mating type following germination, returning to the diploid state (Bai et al. 2022). In the absence of a mating partner, haploid cells undergo mating type switching during mitosis, allowing for mating and diploid formation (“haplo-selfing”) (Hittinger 2013). Wild isolates of this species are generally diploid (Peter et al. 2018), reflecting the characteristics above; this organism is described as “diplontic”, meaning that it displays a strong bias towards growth in the diploid state (Hittinger 2013; Hanson and Wolfe 2017). Observations that other *Saccharomyces* species are also diplontic (Johnson et al. 2004; Lopes et al. 2010; Hittinger 2013; Molinet et al. 2022), imply that the lineage leading to *S. cerevisiae* has experienced selection primarily in the diploid state for at least 14 million years (Shen et al. 2018). Most laboratory strains of *S. cerevisiae* have been modified to lack mating type switching, allowing for the maintenance of asexual haploid populations. A mutation accumulation (MA) study of asexual haploid and diploid *S. cerevisiae* from the same genetic background found that mutations occurred at a higher rate in haploids, particularly in late-replicating regions of the genome (Sharp et al. 2018). This finding is consistent with the idea that selection generally favors reduced mutation rates but has failed to prevent the emergence of mutator alleles with haploid-specific effects, due to the historical rarity of that cell type. Alternatively, this pattern could reflect a higher mutability of haploid genomes, regardless of which cell type is more common.

Here, we sought to clarify the roles of evolutionary history and cell type in shaping mutation patterns by examining the “haplontic” fission yeast *Schizosaccharomyces pombe*. In this species, most growth occurs in asexual haploids; mating between compatible haploid cells is induced by nitrogen limitation, and the resulting diploids sporulate immediately, except in specific lab strains where meiosis is prevented, allowing for the maintenance of vegetative diploids (Sabatinos and Forsburg 2010). We conducted MA in 100 lines of initially isogenic haploid and diploid *S. pombe* carrying the mat2-102 allele, which prevents meiosis in diploids, by imposing 100 single-cell bottlenecks over the course of 210 days. In this species, we find a highly elevated mutation rate in diploid cells––the rare cell type––indicating that a lack of effective selection on rare cell types drives their mutation rates up, as predicted under the drift barrier hypothesis.

## Materials and Methods

### Media and strain preparation

Unless otherwise noted, we conducted our experiments on solid yeast-peptone-dextrose (YPD) media at 30 C. We obtained strain 2172 from the National Collection of Yeast Cultures (Norwich, UK), which is homothallic and carries the mat2-102 allele. This strain can mate with itself but is incapable of meiosis (Willer et al. 1995; Forsburg 2003). In the process of conducting our study we established that the mat2-102 allele is a T-to-G substitution at nucleotide position 449 of mat2-Pi, causing Leu150*, i.e., a premature stop codon, truncating the protein by 10 amino acids and affecting the homeodomain. We isolated a single colony from this strain, grew a liquid culture, spotted on SPAS media, and incubated at 25 C to induce mating (Forsburg and Rhind 2006). We then streaked to single colonies on YES media (Forsburg and Rhind 2006), selected colonies at random, and patched them onto YES + phloxin B, on which diploid cultures adopt a slightly darker coloration (Forsburg and Rhind 2006). We selected darker patches, streaked to single colonies on YPD, and then prepared the resulting cultures for DNA staining with Sytox Green (Thermofisher S7020) and flow cytometry following established protocols (Sabatinos and Forsburg 2009). We used an isolate with confirmed diploid DNA content, along with a corresponding haploid, to initiate our MA lines by streaking to single colonies and selecting 50 colonies of each cell type at random.

### Mutation accumulation

To conduct mutation accumulation, we bottlenecked replicate cultures to single cells every 2-3 days, thereby imposing a low effective population size that prevents selection from acting effectively on most new mutations. We conducted 100 single-cell bottlenecks for each line over the course of 210 days, using sterile toothpicks to transfer the colony closest to the center of each plate. We kept previous plates at 4 C as backups and used these in rare instances where streaking was unsuccessful. Five MA lines were re-initialized from existing MA lines due to bacterial contamination, and this shared history was accounted for in our analyses (one shared mutation was identified and randomly assigned to one of the lines involved). Following MA, we grew the lines in liquid YPD and used these cultures for three purposes: (i) flow cytometry to assess ploidy following the protocol described above, (ii) genomic DNA extraction as described below, and (iii) preservation in 15% glycerol at –80 C.

### Cell counts

We performed cell counts on randomly selected haploid and diploid MA lines throughout the experiment by suspending single colonies in sorbitol and counting cells in a hemocytometer. We estimated cell divisions per unit time assuming a constant rate of doubling as *log*_2_(*N_t_*)/*t*, where *N_t_* is the number of cells in a colony and *t* is the time elapsed since streaking. We estimated the effective population size under MA as the harmonic mean of population sizes in a colony between bottlenecks.

### DNA extraction and sequencing

We extracted genomic DNA from liquid cultures of 100 MA lines and 3 ancestral haploids using the Zymo YeaStar Genomic DNA Kit with the chloroform protocol (Zymo Research, Irvine, CA) and measured DNA concentration using the Qubit dsDNA HS Assay Kit (Life Technologies, Grand Island, NY). Sequencing libraries were prepared by the University of Wisconsin-Madison Biotechnology Center according to the QIAGEN FX DNA Library Preparation Kit (QIAGEN). Quality and quantity of the finished libraries were assessed using an Agilent Tapestation (Agilent, Santa Clara, CA) and Qubit dsDNA HS Assay Kit, respectively. Paired end, 150 bp sequencing was performed using the Illumina NovaSeq6000 (Illumina, San Diego, CA).

### Bioinformatics pipeline

We trimmed reads using *BBDuk* (Energy Joint Genome Institute), mapped reads to the *S. pombe* reference genome ASM294v2 using Burrows Wheeler Aligner *mem* (Li and Durbin 2009), and removed duplicates using Picard Tools (https://broadinstitute.github.io/picard). We identified variants using Haplotype Caller in the Genome Analysis Toolkit (DePristo et al. 2011), applying the appropriate “ploidy” argument for each sample. We excluded variants displaying high strand bias (SOR > 5) or low quality by depth (QD < 2) and considered only sites with coverage in at least 90% of samples, where sequencing depth in the focal sample did not exceed two-times the chromosome-wide median. We used matching criteria to determine the number of callable sites in each sample; applying more stringent calling criteria did not affect our conclusions (SI Appendix). We considered only variants detected in just one MA line; following (Sharp et al. 2018), the probability of missing more than one true mutation in our experiment as a result of this criterion is 0.038%. We used Sanger sequencing to confirm a random subset of putative SNMs, and successfully confirmed 34 out of 35, with no difference in the rate of confirmation between haploids and diploids (Fisher’s exact test: *P* = 1).

## Results

Our results refer to *S. pombe* except where noted; we reiterate published findings for *S. cerevisiae* for comparison (Sharp et al. 2018), and present new analyses of the *S. cerevisiae* data as needed.

### Growth rates and ploidy change

Throughout the experiment, we sampled colonies from MA plates and conducted cell counts to estimate rates of cell division in haploids and diploids (Table 1). We found that the average rate of cell division was about 3% higher in haploids (*t* = 2.47, *df* = 145, *P* = 0.015), with effective population sizes of 12.2 and 11.9 for haploid and diploid MA lines, respectively. Following MA, we assessed the ploidy of each line using flow cytometry; we found that one haploid line had become diploid, and six diploid lines had become haploid, with no significant difference in these frequencies (Fisher’s exact test, *P* = 0.11). Our best estimates of the rate of ploidy change per cell division for this strain are 1.0 × 10^−5^ in haploids and 6.6 × 10^−5^ in diploids. We excluded MA lines that switched ploidy from our analyses, but we note that they did not display unusual mutation rates (Fig. S1).

**Table 1.** Sample sizes, generations, and mutation rates.

|  | Haploid | Diploid |
| --- | --- | --- |
| N lines* | 49 | 44 |
| Mean generations | 1986.9 | 1926.9 |
| Mean coverage | 25.5 | 32.7 |
| Mean callable sites | 12417152 | 12435857 |
| Single-site mutation | $n = 134, \mu = 1.11 (0.93\text{--}1.31) \times 10^{-10}$ | $n = 306, \mu = 1.56 (1.39\text{--}1.74) \times 10^{-10}$ |
| Multi-site mutation | $n = 5, \mu = 0.41 (0.15\text{--}0.89) \times 10^{-11}$ | $n = 69, \mu = 3.51 (2.74\text{--}4.41) \times 10^{-11}$ |
| Insertion mutation | $n = 90, \mu = 7.44 (6.01\text{--}9.09) \times 10^{-11}$ | $n = 160, \mu = 8.14 (6.94\text{--}9.48) \times 10^{-11}$ |
| Deletion mutation | $n = 16, \mu = 1.32 (0.78\text{--}2.08) \times 10^{-11}$ | $n = 69, \mu = 3.51 (2.74\text{--}4.41) \times 10^{-11}$ |

\*Excludes lines that changed ploidy

For each MA line, we examined sequencing coverage on each chromosome to identify cases of possible aneuploidy. We identified two diploid lines with elevated coverage on chromosome III, with median coverage about 1.3 to 1.4 times the median for the other chromosomes (Fig. S2). These elevations are less than the value of 1.5 expected under trisomy, which could indicate that some fraction of the sequenced populations had reverted to euploidy at the time of DNA extraction. These data imply that the rate of mutation to trisomy-III in diploids is at least 2.1 × 10^−5^ per generation.

### Point mutations

Our main goal was to compare the effect of ploidy on the mutation process in the haplontic yeast *S. pombe* (this study) with the pattern in the diplontic yeast *S. cerevisiae* (Sharp et al. 2018). We identified 834 point mutations in total (Table S1), including single nucleotide mutations (SNMs), insertions and deletions (indels), and events that affected multiple nearby sites (each counted as one event for rate analyses). We did not detect any mutations in the mitochondrial genome (upper 95% CI for mitochondrial mutation rate: 1.26 × 10^−9^). We estimated nuclear mutation rates accounting for the number of generations, the number of callable sites, and loss of heterozygosity in diploids. We found that the SNM rate was 60% higher in diploids (Table 1; Fig. 1; likelihood ratio test, LRT: *P* = 1.6 × 10^−6^), in contrast to *S. cerevisiae*, where this rate was 40% higher in haploids (Sharp et al. 2018). We also found that the indel rate was 33% higher in diploids (Table 1; Fig. 1; LRT: *P* = 0.014), in contrast to *S. cerevisiae* where no significant ploidy effect was detected for this mutation type.

**Figure 1.**
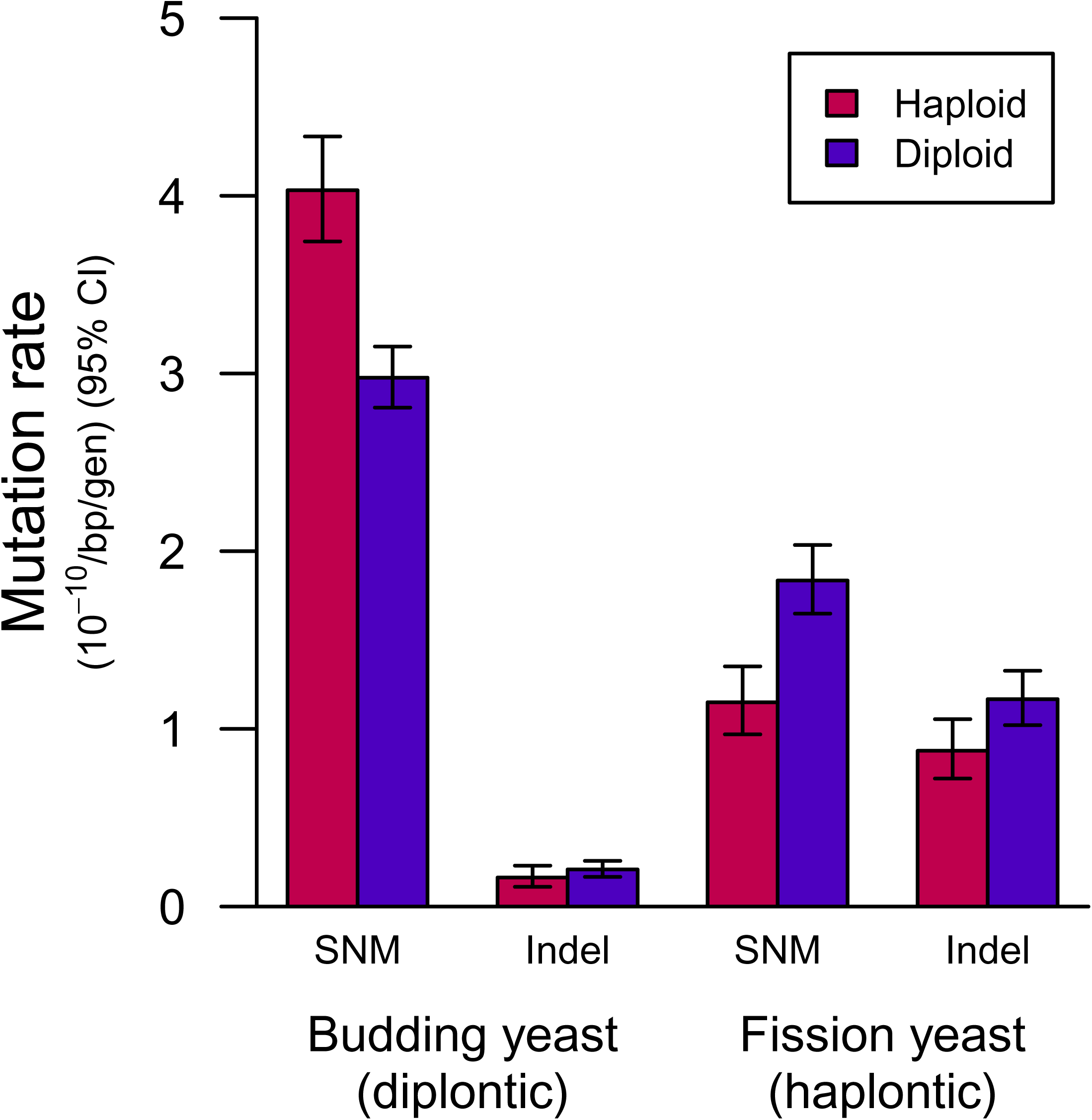
Point mutation rates in budding yeast and fission yeast grown as haploids or diploids. The diplontic budding yeast *S. cerevisiae* shows an elevated rate of single-nucleotide mutations (SNM) in haploid cells and no effect of ploidy on the indel rate. The haplontic fission yeast *S*. *pombe* shows elevated rates of SNM and indel events in diploid cells. Rates are expressed as events per base pair (bp) per generation (gen). Mutation events affecting multiple sites are included in SNM if no indels are present and included in indels otherwise. Error bars represent 95% confidence intervals. Budding yeast data were derived from (Sharp et al. 2018).

In principle, a lower realized rate of mutation accumulation in haploids than in diploids could occur if many new mutations have strongly deleterious recessive effects, making them less likely to accumulate under MA. To study this possibility, we looked for differences in the frequency of mutations with putatively functional consequences in haploids versus diploids (Fig. S3). We found no ploidy difference in the frequency of mutations in coding sequences (χ^2^ = 1.45, *df* = 1, *P* = 0.23). Among mutations in coding sequences, we found no ploidy difference in the frequency of mutations in genes classified as essential (χ^2^ = 0.52, *df* = 1, *P* = 0.47), or in the frequency of synonymous versus nonsynonymous mutations (χ^2^ = 1.91, *df* = 1, *P* = 0.17). Among nonsynonymous mutations, we found no ploidy difference in the frequency of missense versus nonsense mutations (χ^2^ = 0.26, *df* = 1, *P* = 0.61). We therefore see no evidence that selection differentially affected mutation accumulation in haploids and diploids.

### Mutation spectrum

The molecular spectrum and genomic locations of mutations are topics of interest in their own right, but spectrum differences between cell types can also provide insight into the DNA replication and repair processes driving the net difference in mutation rate. We found that the number of mutations per chromosome did not deviate from the expectation based on chromosome length for either cell type (haploids: χ^2^ = 2.02, *df* = 2, *P* = 0.36; diploids: χ^2^ = 0.38, *df* = 2, *P* = 0.83). We examined the rate of mutation events (SNMs or indels) that affected multiple nearby sites within the same MA line, which is a phenomenon that has been observed previously in haploid *S. pombe* (Farlow et al. 2015; Behringer and Hall 2016) and in other species (Schrider et al. 2011). The mutations in diploids were more likely to belong to this category than the mutations in haploids (χ^2^ = 20.03, *df* = 1, *P* = 7.6 × 10^−6^). Analyzing single-site and multi-site events separately, we found that the single-site rate was 1.4-fold higher in diploids (LRT: *P* = 8.7 × 10^−4^, Fig. 2) and that the multi-site rate was 8.5-fold higher in diploids (LRT: *P* = 3.9 × 10^−10^, Fig. 2). In contrast, the frequency of multi-site events did not differ between cell types in *S. cerevisiae* (χ^2^ = 0.06, *df* = 1, *P* = 0.80).

**Figure 2.**
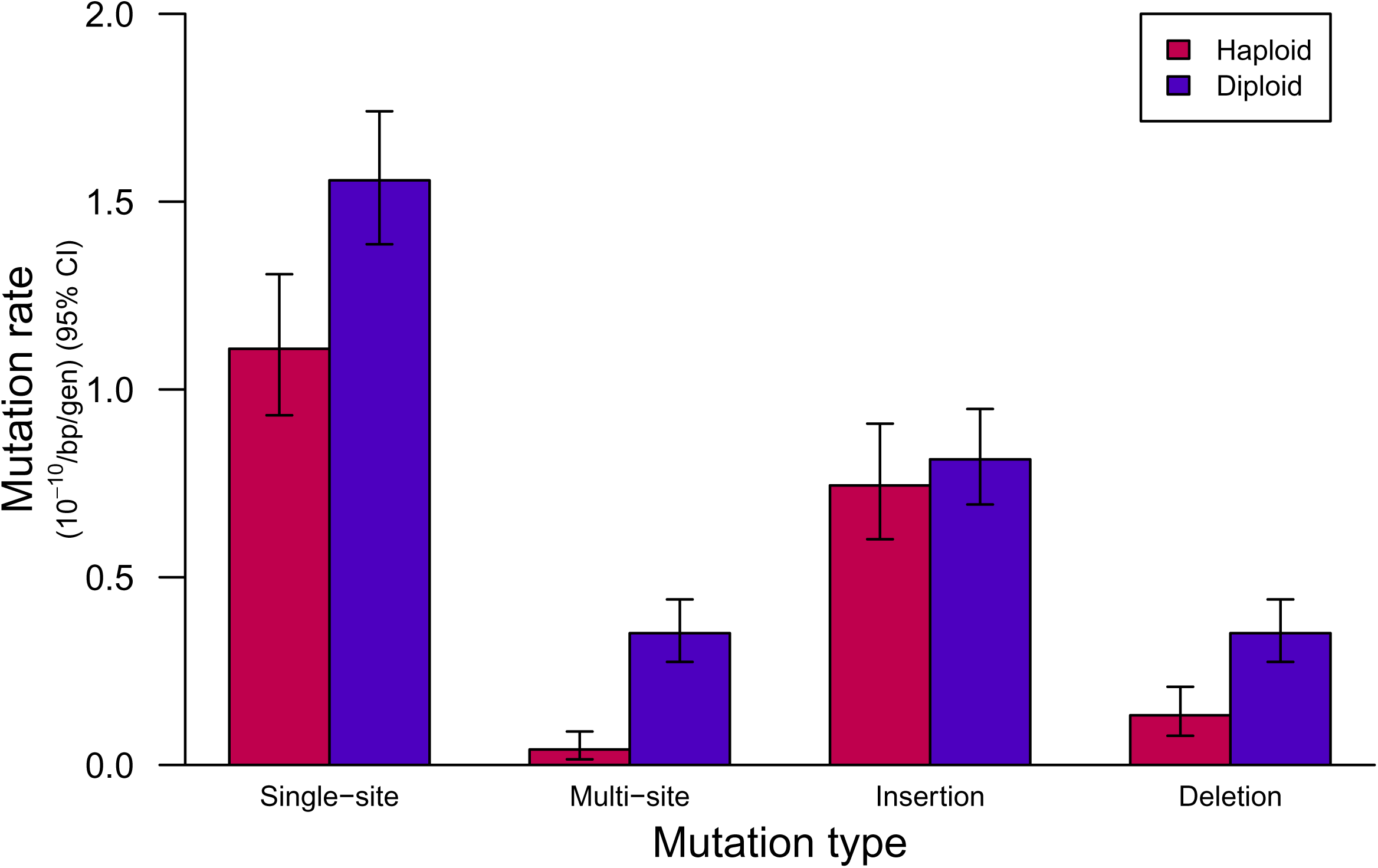
Point mutation spectrum in fission yeast grown as haploids or diploids. Diploids showed elevated rates of single nucleotide changes at single sites and events affecting multiple sites. Ploidy did not affect the rate of insertions, but diploids showed an elevated deletion rate. Error bars represent 95% confidence intervals.

We considered the genomic locations of mutations by defining their relative distance from a centromere. For single-site events, we found no evidence that the locations of mutations differed between cell types (Wilcox test: *P* = 0.40). However, multi-site events differed in location, being farther from centromeres in haploids than in diploids (Fig. 3A; Wilcox test: *P* = 0.019). Using a randomization test, we confirmed that the effect of ploidy on mutation locations differed significantly between multi-site events and single-site events (*P* < 1 × 10^−4^). In other words, multi-site mutation events tended to occur far from centromeres in haploids, whereas in diploids these events were distributed throughout each chromosome (Fig. 3A).

**Figure 3.**
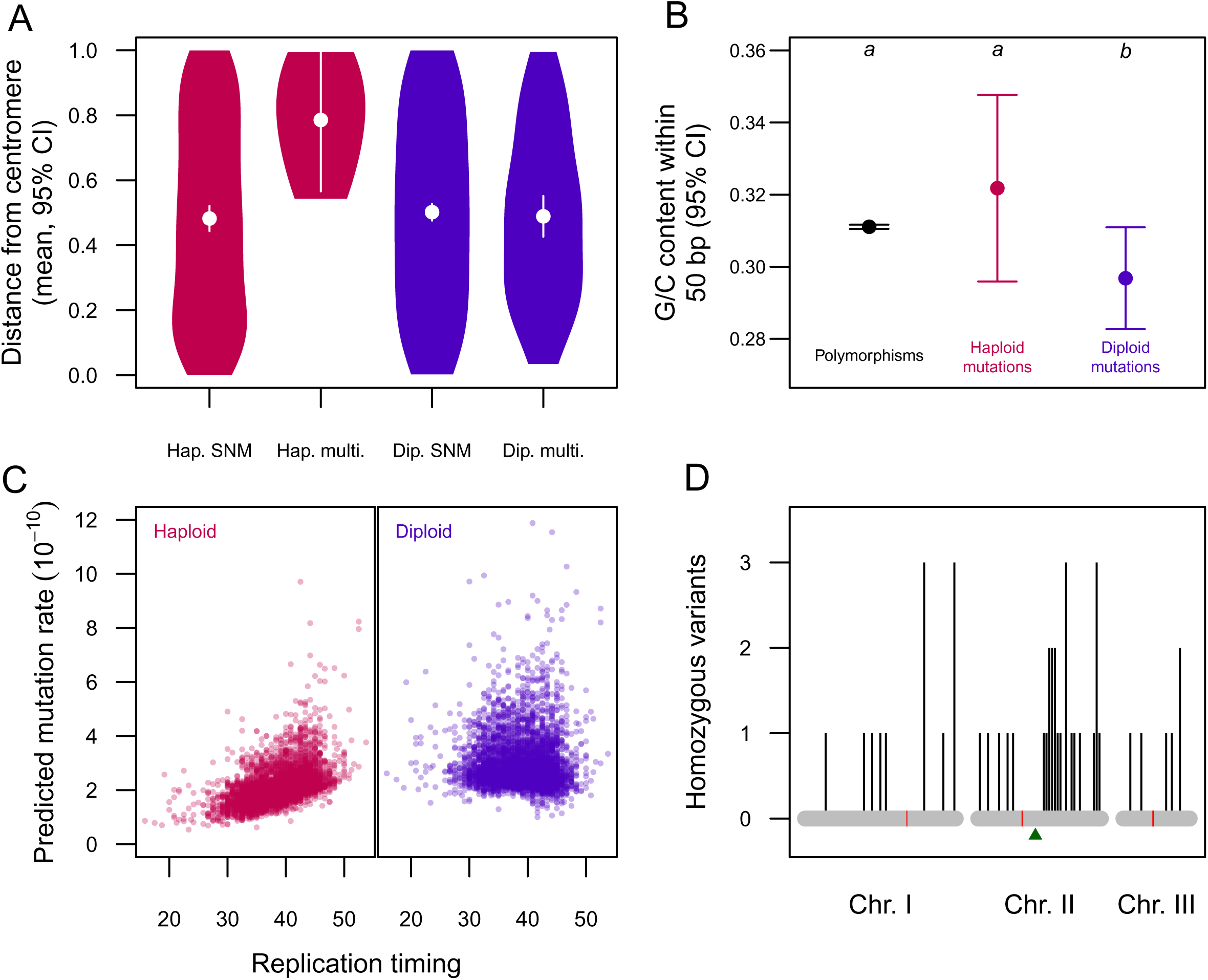
Mutation spectrum in fission yeast grown as haploids or diploids. (A) In haploids, multi-site mutations tended to occur towards the ends of chromosomes, whereas in diploids such event occur throughout chromosomes. (B) The average G/C content within 50 base pairs (bp) of mutations in haploids was not significantly different from that of polymorphisms, whereas mutations in diploids occurred in lower G/C regions, on average. Categories marked with different letters are statistically different. (C) Mutation patterns with respect to replication timing across 4733 genomic windows, as predicted by a generalized linear model accounting for G/C content and detection power in each window. We detected a significant timing-by-ploidy interaction effect, where mutations were more likely in late-replicating regions in haploids but not in diploids. (D) Genomic distribution of homozygous variants in diploids, reflecting loss of heterozygosity. Homozygous variants were more likely on chromosome II; within chromosome II, these variants were more likely to be distal to the mating type locus (green triangle) than expected by chance.

As in previous studies of haploid *S. pombe* (Farlow et al. 2015; Behringer and Hall 2016), we found that indel mutations were biased towards insertions in both cell types (binomial tests; haploids: *P* = 3.5 × 10^−13^; diploids: *P* = 1.9 × 10^−8^). However, the ratio of insertions to deletions differed between haploids and diploids (χ^2^ = 8.65, *df* = 1, *P* = 0.003), and this remained the case when indels that were involved in multi-site events were excluded (χ^2^ = 6.15, *df* = 1, *P* = 0.013). Analyzing insertions and deletions separately, we found that the insertion rate in diploids was not significantly different from the haploid insertion rate (Fig. 2; LRT: *P* = 0.50), whereas the deletion rate was 2.7-fold higher in diploids (Fig. 2; LRT: *P* = 1.2 × 10^−4^). In other words, the higher indel rate we observed in diploids was driven by an increased deletion rate. We found no evidence for differences in length for indels occurring in haploids versus diploids (Wilcox tests; insertions: *P* = 0.46; deletions: *P* = 0.21). In contrast, ploidy did not affect the insertion-deletion ratio in *S. cerevisiae* (χ^2^ = 0.39, *df* = 1, *P* = 0.53).

Another key aspect of the mutation spectrum is the frequency of each SNM type, and non-random distributions have been observed in haploid *S. pombe* (Farlow et al. 2015; Behringer and Hall 2016), haploid and diploid *S. cerevisiae* (Zhu et al. 2014; Sharp et al. 2018; Liu and Zhang 2019), and other organisms. As in *S. cerevisiae* (Sharp et al. 2018), we found that the SNM spectrum differed between haploids and diploids in *S. pombe* (Fig. 4; χ^2^ = 24.45, *df* = 5, *P* = 1.8 × 10^−4^). This was true for mutations at A/T sites (χ^2^ = 13.78, *df* = 2, *P* = 0.001) and G/C sites (χ^2^ = 10.70, *df* = 2, *P* = 0.005); this was also true for single-nucleotide mutations involved in multi-site events (χ^2^ = 12.9, *df* = 5, *P* = 0.024), or not (χ^2^ = 22.0, *df* = 5, *P* = 5.1 × 10^−4^).

**Figure 4.**
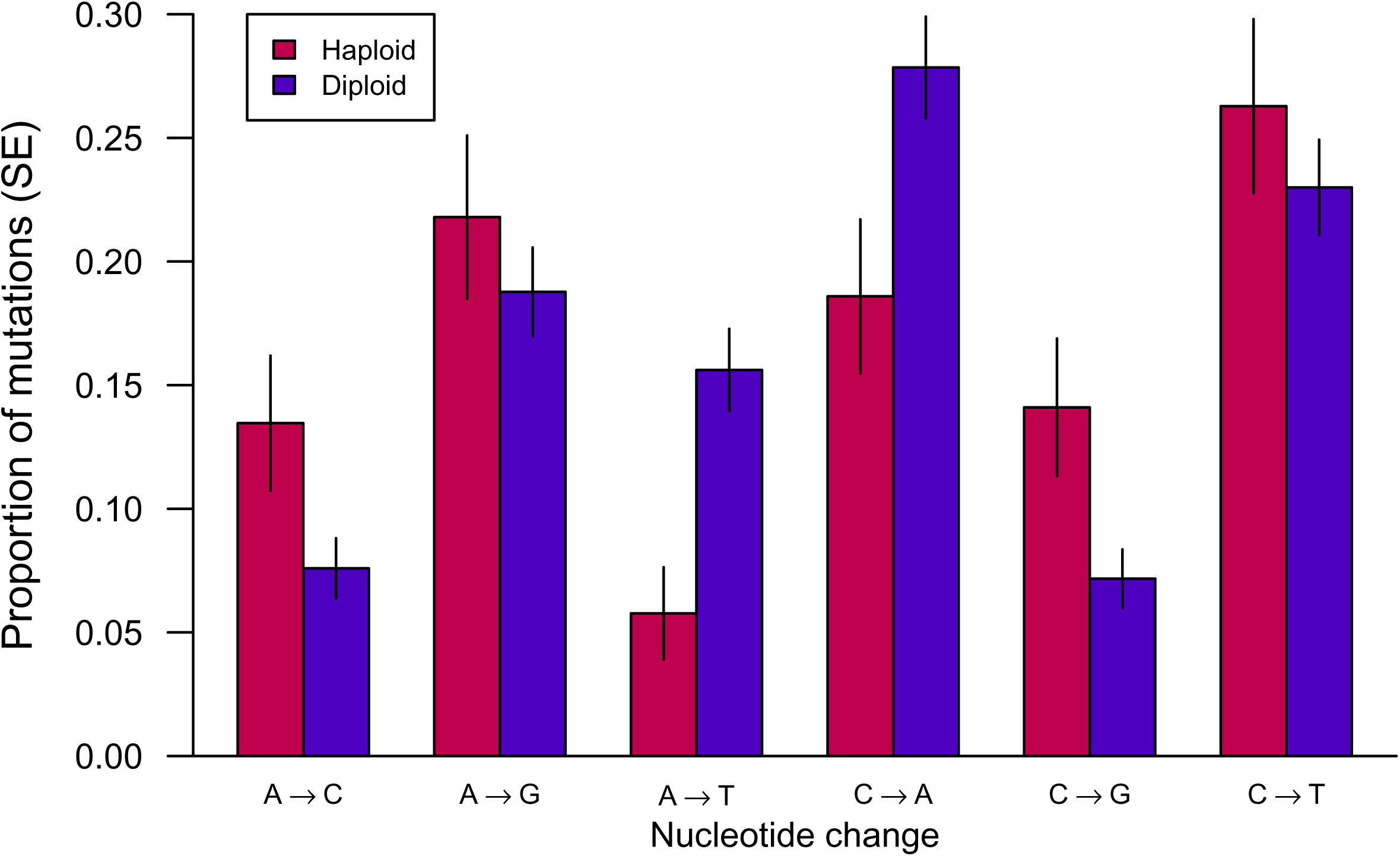
Spectrum of single-nucleotide mutations in fission yeast grown as haploids or diploids. The proportion of mutations across six categories differed between haploids and diploids.

We also considered the nucleotide context in which SNMs occurred, specifically the 3-bp context (where the central base mutated; 32 categories) and the G/C content of the sites 50 bp upstream and downstream of each mutant site, which have both been found to correlate with mutation patterns in previous studies (e.g., (Fryxell and Moon 2005; Sharp and Agrawal 2012; Ness et al. 2015; Kiktev et al. 2018; Sharp et al. 2018)). We found differences in the 3-bp context profile between haploid and diploid mutations outside of protein coding sequences (Table S2; χ^2^ = 46.32, simulated *P* = 0.028), but not within protein coding sequences (Table S2; χ^2^ = 31.51, simulated *P* = 0.44), possibly reflecting limitations on the diversity of nucleotide contexts within coding regions (randomly-generated mutations at callable sites also show a difference in context profile between coding and non-coding regions: χ^2^ = 964.6, *df* = 31, *P* < 2.2 × 10^−16^). Similarly, the average G/C content within 50 bp of mutations was higher in haploids than in diploids (Fig. 3B, *t* = 3.01, *df* = 628, *P* = 0.0027), including when accounting for the coding status of mutations in a linear model (*t* = 2.10, *P* = 0.036). In *S. cerevisiae*, there was little evidence that ploidy affected these aspects of mutation context (Sharp et al. 2018).

In *S. cerevisiae*, ploidy particularly influenced the rate and spectrum of single-nucleotide mutations in late-replicating regions of the genome (Sharp et al. 2018). We examined *S. pombe* mutation in relation to data from (Eshaghi et al. 2007), where replication timing was characterized across 4733 loci in haploids. Our analysis assumes that haploids and diploids have the same replication timing profile, as in *S. cerevisiae* (Müller et al. 2014). Accounting for G/C content and detection power in each bin, we found a significant timing-by-ploidy interaction effect (Fig. 3C; quasi-Poisson GLM: *t* = 2.25, *P* = 0.024; these results include all point mutation types, which showed consistent patterns). Separating haploids and diploids, we found a significant positive effect of replication timing in haploids (Fig. 3C; *t* = 5.58, *P* = 2.5 × 10^−8^) and a non-significant effect in diploids (Fig. 3C; *t* = 1.21, *P* = 0.23). We found no evidence for a difference in the SNM spectrum or insertion-deletion ratio between early-and late-replicating regions for either ploidy type (SNMs: haploids: χ^2^ = 4.57, *df* = 5, *P* = 0.47, diploids: χ^2^ = 4.52, *df* = 5, *P* = 0.48; indels: haploids: χ^2^ = 0.89, *df* = 1, *P* = 0.35, diploids: χ^2^ = 0.26, *df* = 1, *P* = 0.61), in contrast to the pattern observed in *S. cerevisiae* (Sharp et al. 2018).

Since haploidy is the predominant cell type in natural populations of *S. pombe*, we predicted that patterns of polymorphism in such populations would more closely reflect the mutation patterns we observed in haploids, rather than diploids. We considered the polymorphism data from (Jeffares et al. 2015), restricting our analyses to non-genic SNPs with quality scores of at least 30. To compare mutations with the spectrum of polymorphisms where the ancestral base is unknown, we considered the “folded” mutation spectrum with four categories rather than six (Farlow et al. 2015); this differs between haploid and diploid mutations (χ^2^ = 16.4, *df* = 3, *P* = 9.2 × 10^−4^). The mutation and polymorphism spectra differ for haploids (χ^2^ = 66.0, *df* = 3, *P* = 3.1 × 10^−14^), as observed in a previous study (Farlow et al. 2015), and for diploids (χ^2^ = 144.1, *df* = 3, *P* < 2.2 × 10^−16^). It is therefore not clear whether polymorphisms more closely reflect the haploid mutation process in this regard. However, we found that polymorphisms were more like haploid mutations in three other respects. First, the 3-bp context profile of non-coding polymorphisms was not statistically distinguishable from the profile of non-coding mutations in haploids (Table S2; χ^2^ = 40.32, simulated *P* = 0.14) but was significantly different from the diploid profile (Table S2; χ^2^ = 53.74, simulated *P* = 0.004). Second, the average G/C content surrounding non-coding polymorphisms differed from that of non-coding diploid mutations (Fig. 3B; Welch *t* = 1.99, *df* = 168.58, *P* = 0.048), but not from that of haploid mutations (Fig. 3B; Welch *t* = –0.84, *df* = 37.04, *P* = 0.41). Third, we found that polymorphisms were significantly more common in later-replicating regions (linear model: *t* = 2.26, *P* = 0.0236), matching the pattern observed in haploids but not diploids. To summarize, there are several aspects of the mutation spectrum that differ between haploids and diploids, where the haploid mutation pattern appears to be reflected in the genomic distribution of polymorphisms.

### Loss of heterozygosity

In the absence of sexual reproduction, new mutations in diploids can still become homozygous through homologous repair of DNA double strand breaks. Such loss-of-heterozygosity (LOH) events have been observed in asexual diploid *S. cerevisiae* (e.g., (Andersen et al. 2008; Yim et al. 2014; Sharp et al. 2018; Nguyen et al. 2020; Dutta et al. 2022; Tutaj et al. 2022; Vijayan et al. 2025)), and in screening assays with diploid *S. pombe* (Cullen et al. 2007; Park et al. 2025). In our study, we identified 44 independent LOH events across 23 diploid MA lines. Following the approach in (Sharp et al. 2018), our mutation rate estimation procedure accounted for mutations in diploids that became unobservable due to LOH towards the ancestral allele and allowed us to jointly estimate the rate of LOH. We found that LOH in *S. pombe* occurred at a rate of 1.6 × 10^−4^ per diploid site per generation (95% CI: 1.1 to 2.1 × 10^−4^), which is 2.0 times higher than the LOH rate observed in *S. cerevisiae* (95% CI: 1.3 to 3.1). Among diploid mutations, we found that the frequency of homozygous mutations relative to heterozygous mutations was elevated on chromosome II (Fig. 3D; χ^2^ = 9.96, *df* = 2, *P* = 0.007). The distribution of variants in terms of distance from a centromere differed between homozygous and heterozygous variants (KS test, *P* = 0.015), with homozygous variants further from centromeres on average (Welch *t* = 2.05, *df* = 53.0, *P* = 0.046), but these effects did not persist if chromosome II was excluded. The frequency of homozygous variants did not differ between the left and right arm of chromosome II (Fig. 3D; binomial test: *P* = 0.10); however, chromosome II also contains the mating type locus, and we found that 21 of the 26 homozygous variants on this chromosome were distal to this locus, more than expected by chance (Fig. 3D; binomial test: *P* = 0.005), suggesting that the DNA breaks at this locus that give rise to mating type switching, including in diploids (Arcangioli and Gangloff 2023), may increase the likelihood of LOH.

## Discussion

We compared ploidy-specific mutation patterns in two species of yeast that differ in life history, with the haploid state predominating in *S. pombe* and the diploid state predominating in *S. cerevisiae*, allowing us to distinguish the effect of ploidy *per se* from the effect of evolutionary history. Our results indicate that there is not a ploidy state that is inherently mutagenic; rather, mutation rates are elevated in whichever cell type is less common (Fig. 1), as predicted under the drift barrier hypothesis. A possible exception is aneuploidy events, which we observe in diploids and not haploids of both species, though this could reflect stronger selection against aneuploidy in haploids. If mutation rates were instead under stabilizing selection, the mutation rate in rare cell types would be expected to drift in either direction, depending on the net combination of mutator and anti-mutator alleles specific to that cell type.

We find that haploid and diploid *S. pombe* also differ in multiple aspects of the mutation spectrum, with diploids displaying a shift towards insertions and multi-site mutation events (Fig. 2). The genomic distribution of mutations also differed, with multi-site mutations biased towards the ends of chromosomes in haploids but not diploids (Fig. 3A), and point mutations biased towards late-replicating regions in haploids but not diploids (Fig. 3C). Combined with data on transcription patterns in haploid and diploid *S. pombe* (Park and Forsburg 2024), these patterns may allow the genetic pathways responsible for ploidy differences in mutation to be identified. Ploidy affected the mutation spectrum differently in *S. pombe* than in *S. cerevisiae* (Sharp et al. 2018); given that these species are very distantly related and display different mutation patterns in their dominant cell types, it is perhaps unsurprising that mutation patterns in rare cell types would drift in different ways. We studied mutation spectra in a single genetic background of each species, but in both cases, there is correspondence between mutation patterns in the dominant ploidy type and patterns of polymorphism, indicating that these strains are reasonably representative of natural populations with respect to mutation. The ploidy effect on mutation was associated with replication timing in *S. cerevisiae*, implicating translesion synthesis repair (Sharp et al. 2018); forthcoming work from our lab tests the hypothesis that abolishing translesion synthesis eliminates the ploidy effect in this organism, and a similar approach connecting mutation patterns with repair mechanisms may prove fruitful in *S. pombe*.

Our findings contrasting two species suggest that which ploidy type is historically dominant may have a greater impact on mutation rates than ploidy *per se*. Additional observations in *S. cerevisiae* are consistent with this pattern. Ploidy could directly affect mutation if diploids have greater access to homology directed DNA repair by virtue of having at least two copies of each chromosome present throughout the cell cycle. To test this idea, Sharp et al. (2018) removed the gene *RDH54* from a subset of lines, which is believed to be essential for mitotic recombinational repair between homologous chromosomes in diploids but not between sister chromatids (Klein 1997; McKinney et al. 2013). This gene knockout affected the rate of chromosome segregation errors but had no detectable effect on the point mutation rate (Sharp et al. 2018), suggesting that this type of diploid-specific repair is rare relative to other sources of mutation. Additionally, the ploidy difference in mutation was even stronger in the mitochondrial genome than in the nuclear genome (Sharp et al. 2018), matching other observations (Sia et al. 2000, 2003), even though the mitochondrial genome is typically present in many copies in cells of both types (Grimes et al. 1974; Birky et al. 1978). We did not detect mitochondrial mutations in *S. pombe*, and none were reported in the previous MA experiments with haploids (Farlow et al. 2015; Behringer and Hall 2016). The (callable) size of the mitochondrial genome in *S. pombe* is 26.6% that of *S. cerevisiae*; if mitochondrial point mutations occurred in haploid *S. pombe* at the same rate as diploid *S. cerevisiae*, we would observe no events 18% of the time; further, if the lower overall mutation rate in *S. pombe* (Fig. 1) applies to mitochondria, we would observe no events 33% of the time. Our results are therefore consistent with reasonable mitochondrial mutation rates, but testing for a ploidy effect in this genome would require greater sample sizes. Finally, haploid *S. pombe* spend much of the cell cycle in G2 (Hayles and Nurse 2018), such that templates for homologous repair are usually present (Ferreira and Cooper 2004); diploid also readily undergo homologous repair in mitotic cells, as evidenced by their high rate of LOH (Results; (Park et al. 2025). We suggest that the history of selection under alternative ploidy states may be more relevant to ploidy-specific mutation rate evolution than the mechanics of DNA repair, but evidence from additional taxa would be valuable.

An alternative interpretation of ploidy differences in mutation rate is elevated mutation in the uncommon ploidy type is the result of “stress”. Stress induced mutation can occur in eukaryotes (Agrawal and Wang 2008; Matsuba et al. 2012; Sharp and Agrawal 2012; Shor et al. 2013; Jiang et al. 2014; Liu and Zhang 2021b), though this need not be adaptive, and could itself reflect less effective selection on rarely used DNA repair mechanisms. However, it is not obvious that the rare cell types were stressed in our experiments, as they experienced growth rate reductions of <1% and 3% in *S. cerevisiae* and *S. pombe*, respectively, relative to the common cell type.

In *S. cerevisiae* strains where mating type switching is prevented, diploids can nevertheless arise and spread in haploid populations (Gerstein et al. 2006; Lynch et al. 2008; Venkataram et al. 2016; Fisher et al. 2018; Harari et al. 2018; Gerstein and Sharp 2021), with the inverse scenario occurring less commonly (Zeyl et al. 2003; Crandall et al. 2023). We found that this type of ploidy change can also occur in *S. pombe*, in either direction, at rates similar to estimates from *S. cerevisiae* (Harari et al. 2018; Sharp et al. 2018). In our experiment, the mat2-102 allele (a single-nucleotide substitution causing a premature stop codon; see Materials and Methods) prevents sporulation in diploids (Forsburg 2003); our sequence data confirm that this allele did not revert but stop-codon readthrough might allow for expression in rare instances. Alternatively, transitions from diploidy to haploidy could reflect nondisjunction and genome instability (Bodi et al. 1991; Park et al. 2025). The single case we observed of a haploid line becoming diploid could reflect a rare mating event on rich media, or endoreduplication (Harari et al. 2018) ; this line displays the highest frequency of homozygous variants we observed (40%), consistent with its apparent history.

The yeast species we studied can be grown as asexual haploids or diploids, and this flexibility has allowed for a great deal of valuable genetics research, with haploid *S. cerevisiae* used regularly for convenience. However, our findings suggest that historically rare cell types can exhibit suboptimal patterns of DNA replication and repair, relative to the common cell type, as a result of ineffective purifying selection in the past; in principle, this could apply to other traits as well, adding bias to measures of traits under directional selection and variance to traits under stabilizing selection. Phenotype drift in rare cell types should be limited when there is either a strong genetic correlation for the value of a trait between the rare and common cell type, or strong selection in the rare cell type, e.g., selection on viability or mating competency in haploids. Treating haploids and diploids as interchangeable appears to be unjustified, at least with respect to DNA replication and repair, and we recommend greater consideration of ploidy as a biological variable in genetic studies.

## Data availability

Raw sequencing reads are deposited in the National Center for Biotechnology Information Sequence Read Archive, https://www.ncbi.nlm.nih, accession no. PRJNA1306532. Point mutations identified in *S. pombe* are listed in Table S1.

## Acknowledgements and funding sources

Thanks to Denise Smith for assistance with DNA extraction and Sanger sequencing. Research reported in this publication was supported by the National Institute of General Medical Sciences of the National Institutes of Health under award number R35GM154954 to NPS. The funders had no role in study design, data collection and analysis, decision to publish, or preparation of the manuscript. The authors utilized the University of Wisconsin-Madison Biotechnology Center’s DNA Sequencing Facility (RRID:SCR_017759) and the University of Wisconsin Carbone Cancer Center Flow Cytometry Laboratory, supported by P30 CA014520 from the National Cancer Institute of the National Institutes of Health.

## Supplementary material

### Testing alternative approaches to mutation rate comparisons

Our results suggest that selection did not bias the accumulation of mutations in haploids versus diploids (Fig. S3), but we nevertheless tested for mutation rate differences using only non-protein-coding sites, which are less likely to have experienced effective selection. Repeating our mutation rate analyses using only (callable) non-coding sites, we find essentially the same result as when we included all callable sites: the estimated SNM rate was 2.13-fold higher in diploids (LRT: *P* = 2.10 × 10^−5^), and the estimated indel rate was 1.38-fold higher in diploids (LRT: *P* = 1.87 × 10^−3^). The higher mutation rate in diploids therefore persists when coding regions are excluded.

We also considered whether additional stringency in variant filtering would alter our key results. In addition to the filtering steps described in the Materials and Methods, we removed any variants with Fisher strand bias values of 60 or greater, root mean square mapping quality values less than 40, map quality rank sum values less than –12.5, or read position rank sum values less than –8, and required that at least 90% of alleles at a site were called. Altogether, these steps removed 70 variants, about 6% of the variants originally included. We confirmed that all of our results with respect to mutation rate differences between haploids and diploids remained statistically significant when these additional filters were applied.

Our main analysis approach treats mutation events as independent across MA lines and incorporates a correction for LOH. As a supplementary analysis, we modeled the number of point mutations in each line using a Poisson generalized linear model, with ploidy as a predictor and detection power as a covariate. We expect this approach to underestimate diploid mutation rates by failing to account for LOH. Nevertheless, this model confirmed our finding of a higher mutation rate in diploids (*z* = 4.51, *P* = 6.58 × 10^−6^). Overdispersion was not formally significant (*z* = 1.54, *P* = 0.062), but applying a quasibinomial model did not alter this finding (*z* = 3.61, *P* = 4.97 × 10^−4^). We therefore conclude that our main result is robust to these aspects of the analysis approach.

## Supplementary figure legends

**Figure S1.**
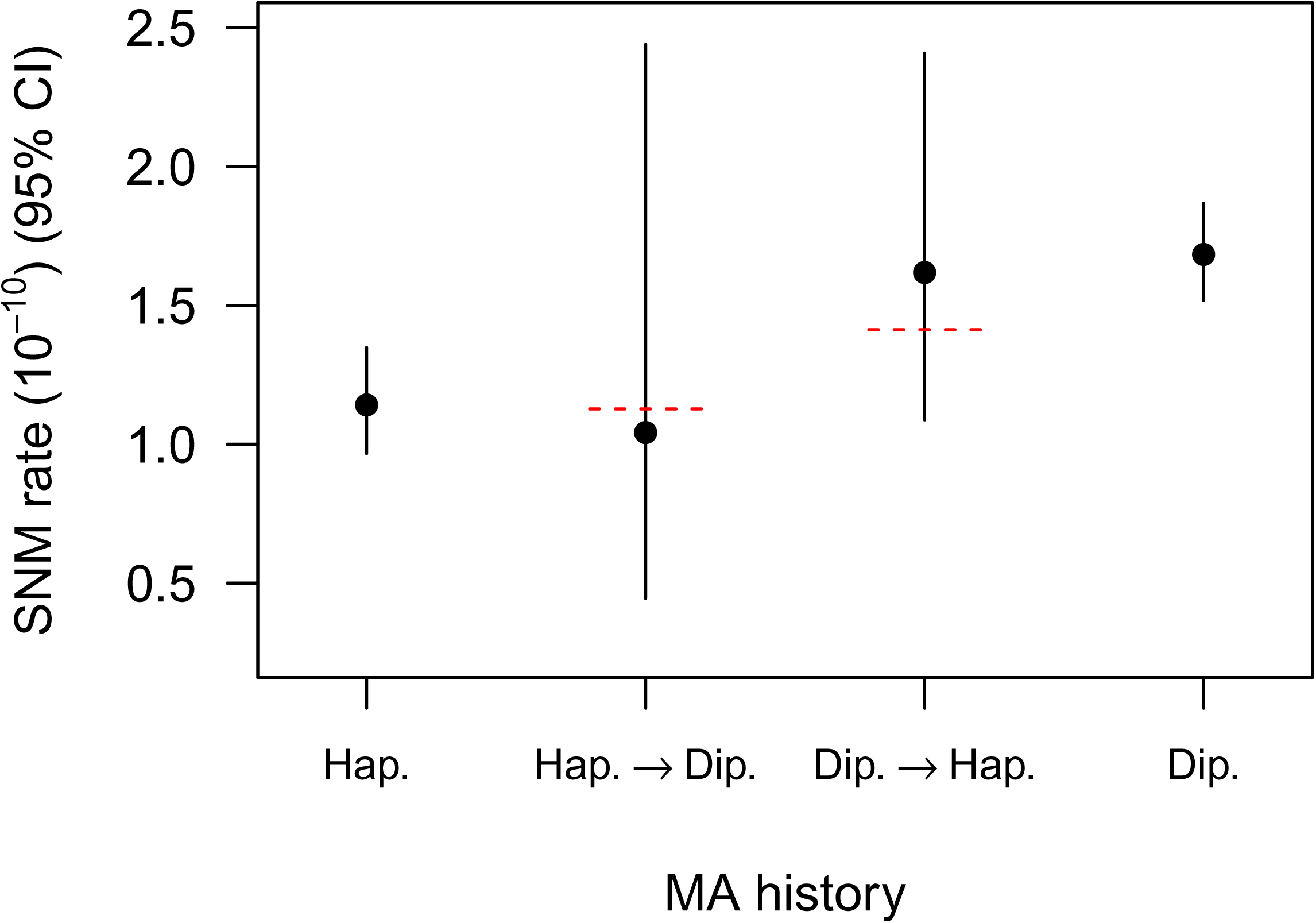
Mutation rates in lines that switched ploidy during mutation accumulation. Rates for lines that did not change ploidy are included for comparison. The apparent mutation rate in lines that switched from haploidy to diploidy is 0.25 µ_haploid_ + 0.5 µ_diploid_, because two homologous chromosomes are scored but only one was present for the haploid phase of the MA line history (half of the experiment duration, on average); since the diploid mutation rate we observed was roughly 1.5-fold that of haploids, this equates to approximately the haploid mutation rate (red dashed line). The apparent mutation rate in lines that switched from diploidy to haploidy is simply the average of the haploid and diploid rates (red dashed line). In both cases, the apparent mutation rates did not differ significantly from the expected rate, though the sample size of lines that switched ploidy is low.

**Figure S2.**
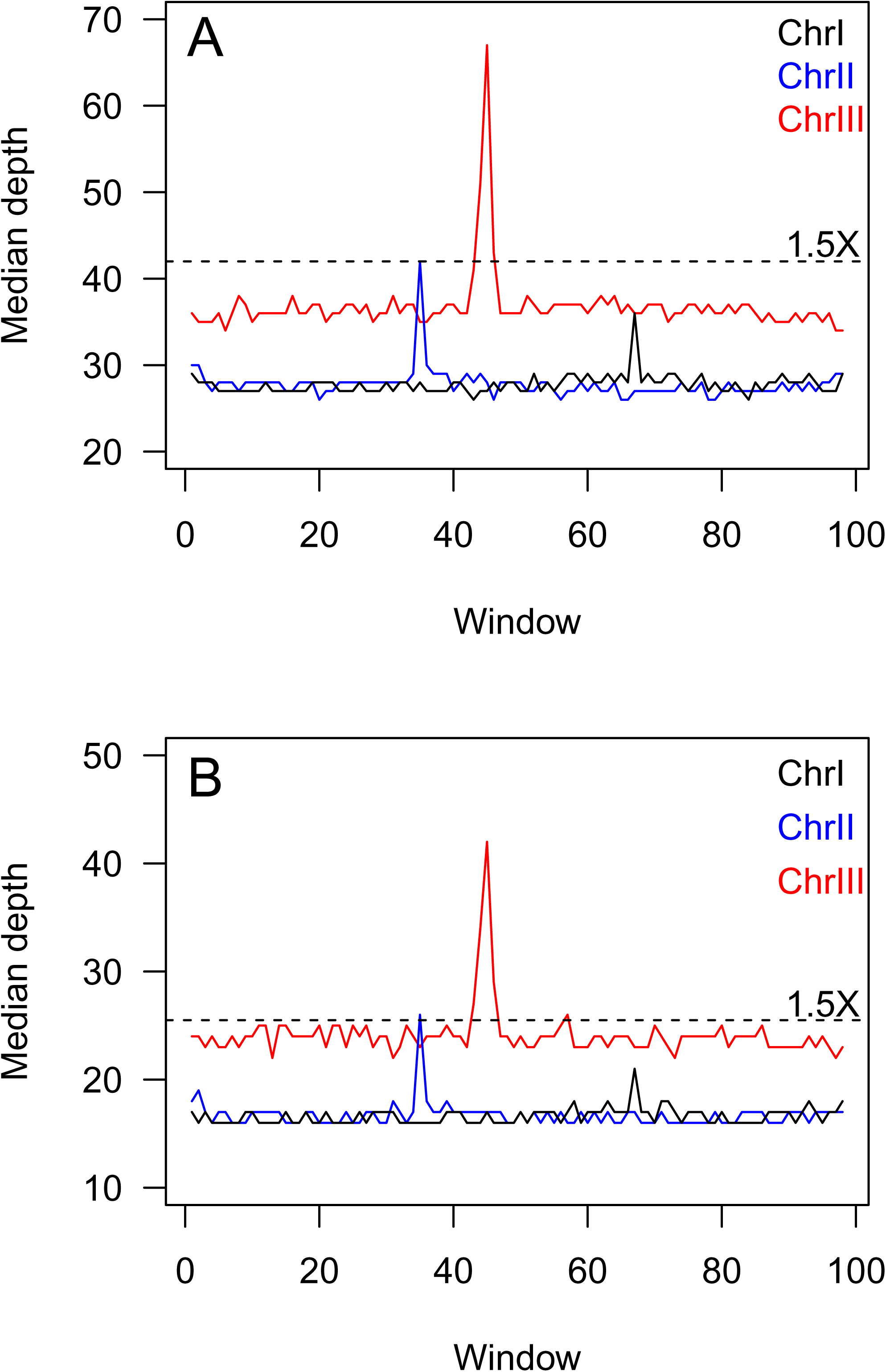
Coverage profiles for each chromosome in two diploid MA lines with evidence of aneuploidy. Lines show median depth in 100 windows across each chromosome, exclude the left- and right-most windows, which are high-coverage regions. Trisomy for a given chromosome is expected to result in 1.5X coverage relative to the other chromosomes (dashed line). These putative cases of aneuploidy, both involving chromosome III, are seemingly not fixed in the cell culture that was sequenced (red line < dashed line), which may indicate that reversion to euploidy was in progress in the sequenced culture. (A) MA line 68. (B) MA line 98.

**Figure S3.**
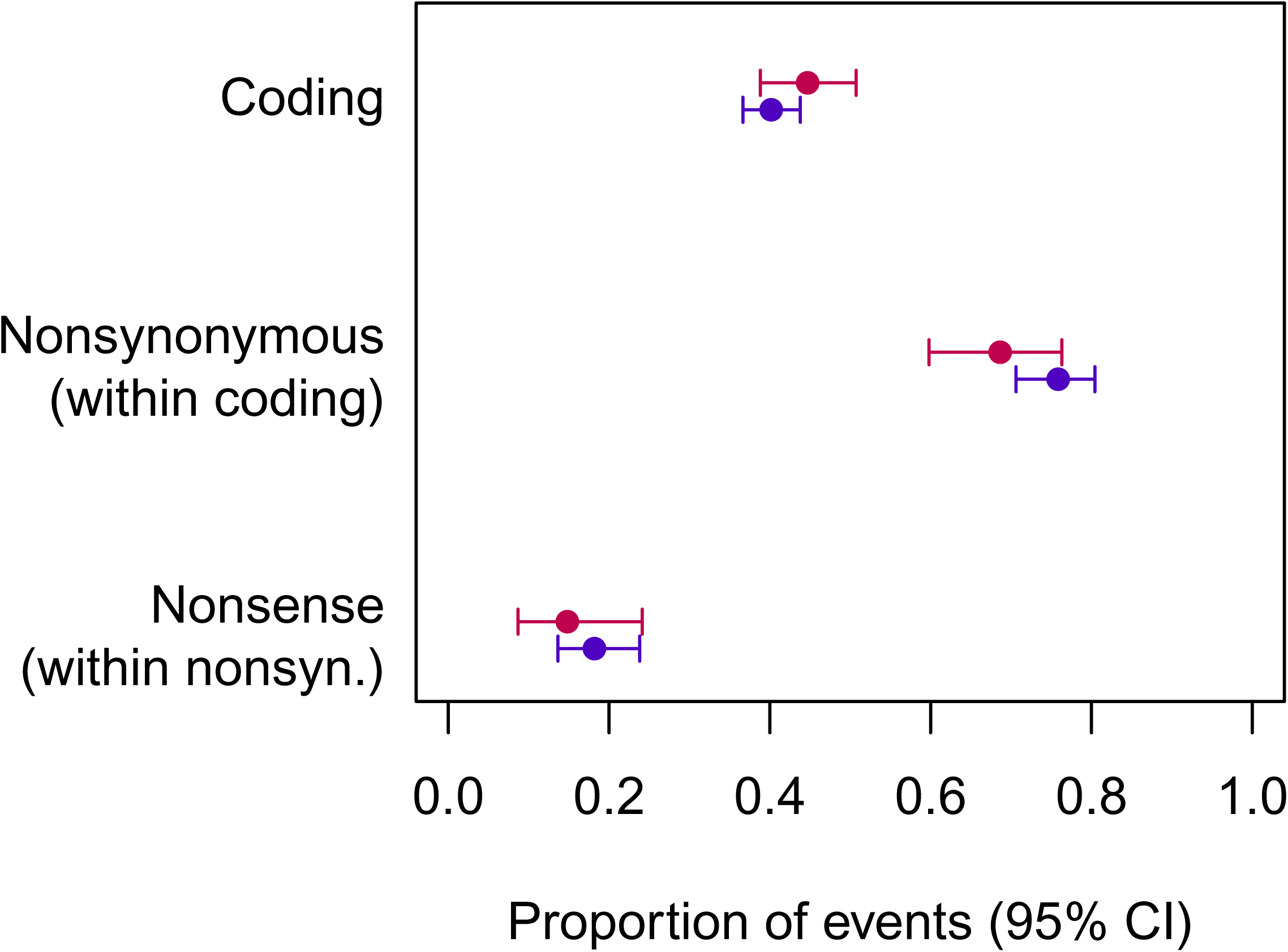
Proportion of mutation events in haploids (red) and diploids (blue) with respect to genic impact. Ploidy did not influence the likelihood that mutations occurred in protein coding regions, not the likelihood that mutations in coding regions were nonsynonymous or nonsense.

## Supplementary tables

**Table S1. Point mutations identified in *S. pombe.* See associated Excel file.**

**Table S2.** Nucleotide context of mutations and polymorphisms.

| Context | All variants |  |  | Non-coding variants |  |  |
| --- | --- | --- | --- | --- | --- | --- |
|  | Haploid | Diploid | Polymorphism | Haploid | Diploid | Polymorphism |
| AAA | 9 | 25 | 7767 | 1 | 14 | 3226 |
| AAC | 4 | 12 | 4364 | 1 | 4 | 1505 |
| AAG | 2 | 10 | 3822 | 0 | 3 | 1270 |
| AAT | 6 | 19 | 8093 | 2 | 4 | 3510 |
| CAA | 4 | 15 | 4798 | 2 | 4 | 1549 |
| CAC | 1 | 9 | 2548 | 1 | 3 | 691 |
| CAG | 1 | 7 | 2715 | 0 | 2 | 566 |
| CAT | 4 | 6 | 4147 | 0 | 3 | 1323 |
| GAA | 5 | 11 | 4113 | 1 | 4 | 1377 |
| GAC | 4 | 8 | 1956 | 2 | 0 | 566 |
| GAG | 1 | 9 | 1945 | 0 | 2 | 519 |
| GAT | 5 | 4 | 3217 | 2 | 0 | 1134 |
| TAA | 6 | 23 | 7385 | 0 | 9 | 3161 |
| TAC | 3 | 8 | 3555 | 1 | 4 | 1371 |
| TAG | 3 | 10 | 3417 | 1 | 6 | 1109 |
| TAT | 6 | 23 | 7456 | 3 | 11 | 3217 |
| ACA | 10 | 33 | 5924 | 3 | 16 | 2383 |
| ACC | 5 | 18 | 3447 | 0 | 3 | 1194 |
| ACG | 8 | 11 | 3601 | 2 | 1 | 1048 |
| ACT | 3 | 25 | 6228 | 1 | 9 | 2429 |
| CCA | 4 | 20 | 3290 | 0 | 5 | 990 |
| CCC | 1 | 5 | 1753 | 0 | 1 | 488 |
| CCG | 6 | 7 | 2356 | 1 | 3 | 495 |
| CCT | 3 | 11 | 3297 | 1 | 5 | 1012 |
| GCA | 14 | 24 | 3538 | 2 | 1 | 1191 |
| GCC | 3 | 7 | 1790 | 2 | 2 | 494 |
| GCG | 7 | 13 | 2242 | 2 | 7 | 618 |
| GCT | 5 | 11 | 3722 | 2 | 4 | 1216 |
| TCA | 6 | 26 | 5674 | 2 | 9 | 2206 |
| TCC | 3 | 20 | 3677 | 0 | 4 | 1191 |
| TCG | 9 | 12 | 3837 | 3 | 9 | 1127 |
| TCT | 5 | 32 | 6423 | 0 | 17 | 2430 |

## Notes

### Competing Interest Statement

The authors have declared no competing interest.

